# Facets of individual-specific health signatures determined from longitudinal plasma proteome profiling

**DOI:** 10.1101/2020.03.13.988683

**Authors:** Tea Dodig-Crnković, Mun-Gwan Hong, Cecilia Engel Thomas, Ragna S. Häussler, Annika Bendes, Matilda Dale, Fredrik Edfors, Björn Forsström, Patrik K.E. Magnusson, Ina Schuppe-Koistinen, Jacob Odeberg, Linn Fagerberg, Anders Gummesson, Göran Bergström, Mathias Uhlén, Jochen M Schwenk

**Affiliations:** Science for Life Laboratory, Department of Protein Science, KTH-Royal Institute of Technology, Tomtebodavägen 23, 171 65 Stockholm, Sweden; Department of Genetics, Stanford University School of Medicine, Stanford, CA 94305, USA; Department of Medical Epidemiology and Biostatistics, Karolinska Institutet, Nobels väg 12A, 171 77 Stockholm, Sweden; Center for Translational Microbiome Research, Department of Microbiology, Tumor and Cell Biology, Karolinska Institute, 171 77 Stockholm, Sweden; K.G. Jebsen Thrombosis Research and Expertise Center (TREC), Department of Clinical Medicine, UiT the Arctic University of Norway, 9010 Tromsø, Norway; Coagulation unit, Department of Hematology, Karolinska University Hospital, 171 76 Stockholm Sweden; Department of Molecular and Clinical Medicine, Institute of Medicine, Sahlgrenska Academy, Gothenburg University, 413 45 Gothenburg, Sweden; Region Västra Götaland, Sahlgrenska University Hospital, Department of Clinical Genetics and Genomics, 413 45 Gothenburg, Sweden; Region Västra Götaland, Sahlgrenska University Hospital, Department of Clinical Physiology, 413 45 Gothenburg, Sweden; Novo Nordisk Foundation Center for Biosustainability, Technical University of Denmark, 2800 Lyngby, Denmark

**Keywords:** affinity proteomics, longitudinal profiling, plasma proteomics, pQTLs, precision medicine

## Abstract

**Background:** Precision medicine approaches aim to tackle diseases on an individual level through molecular profiling. Despite the growing knowledge about diseases and the reported diversity of molecular phenotypes, the descriptions of human health on an individual level have been far less elaborate.

**Methods:** To provide insights into the longitudinal protein signatures of well-being, we profiled blood plasma collected over one year from 101 clinically healthy individuals using multiplexed antibody assays. After applying an antibody validation scheme, we utilized > 700 protein profiles for in-depth analyses of the individuals’ short-term health trajectories.

**Findings:** We found signatures of circulating proteomes to be highly individual-specific. Considering technical and longitudinal variability, we observed both stable and fluctuating proteins in the circulation, as well as networks of proteins that covaried over time. For each participant, there were unique protein profiles and some of these could be explained by associations to genetic variants.

**Interpretation:** This study demonstrates that there was noticeable diversity among clinically healthy subjects, and facets of individual-specific signatures emerged by monitoring the variability of the circulating proteomes over time. Longitudinal profiling of circulating proteomes has the potential to enable a more personal hence precise assessment of health states, and thereby provide a valuable component of precision medicine approaches.

**Funding:** This work was supported by the Erling Persson Foundation for the KTH Centre for Precision Medicine and the Swedish Heart and Lung Foundation for the SCAPIS project. We also acknowledge the Knut and Alice Wallenberg Foundation for funding the Human Protein Atlas project, Science for Life Laboratory for Plasma Profiling Facility, and the Swedish Research Council (Grant no 2017-00641).

## Introduction

Human blood serves as a minimally-invasive source to gain insights about different physiological processes by studying the transcriptome, proteome, or metabolome. Just recently multi-omics studies have emerged to also determine longitudinal profiles of human health and disease [1-3]. Regular monitoring of molecular markers holds the promise to identify perturbations affecting an individual’s baseline levels and follow these changes as a healthy system transitions into a disease state [4]. However, longitudinal studies of clinically healthy subjects remain sparse and limited to certain technologies and analytes.

The blood proteome, consisting of both cellular and soluble proteins, has received a revived interest due to advances in protein technologies. This includes mass cytometry [5] to study immune systems, as well as mass spectrometry [6] and affinity assays [7] for profiling serum or plasma. For the circulating plasma proteome, nearly 5,000 proteins have this far been detected when combing discoveries from all assays and technologies [8]. Surprisingly, only 730 proteins are predicted to be actively secreted into the circulation [9], attributing many of the currently detected proteins to cellular leakage that may possibly occur during sample preparation [10]. Even though highly multiplexed assays have enabled large scale assessment of pre-symptomatic health states [3], additional investigations will complement our molecular description and understanding of the facets of health in healthy individuals.

Here, we used an affinity-based proteomics approach [11] to explore the longitudinal profiles of circulating proteins from 101 clinically healthy individuals selected from the Swedish SCAPIS cohort [12], who donated blood four times during one year. The objective of this study was to capture the signatures and variability of personal plasma proteomes at baseline and follow these proteins over one year.

## Research in context

### Evidence before this study

Proteins circulate the human blood and their analysis can provide important information about health or disease states of an individual. Today, many studies focus on finding proteins related to diseases or specific conditions even though our knowledge about if and why proteins differ between individuals, which protein levels vary over time, and how protein profiles appear in clinically-healthy persons is still limited.

### Added value of this study

Here, we used multiplexed immunoassays to study a large number proteins circulating in plasma of 101 clinically healthy and well characterized individuals over one year. We found a substantial individuality in the protein profiles between the participants, which for some of the proteins could be explained by genetic variants. Our analysis also showed that protein profiles varied among the participants over time, which indicated that a variety of short-term as well as continuous changes can occur even in healthy people.

### Implications of all the available evidence

Our findings add to the understanding of molecular signatures of human health and provide important information for studies aiming at finding common protein biomarkers for diseases. Together with evidence from other studies, it appears necessary to consider the diversity, individuality, and variability over time as critical aspects of molecular signatures that aid to advance precision medicine.

## Materials and methods

### Wellness samples

The Swedish SciLifeLab SCAPIS Wellness Profiling (S3WP) program consists of 101 individuals recruited from the Swedish CArdioPulmonary bioImage Study (SCAPIS), an ongoing prospective observational study [12]. SCAPIS includes 30,154 individuals between 50-65 years that have been randomly selected from the general Swedish population and invited to join the study. All individuals are extensively phenotyped prior to entering the S3WP program. The S3WP study is non-interventional and observational with the aim of collecting longitudinal clinical traits and molecular omics data for all 101 participants. Primary exclusion criteria in SW3P are; 1) previously received health care for myocardial infarction, stroke, peripheral artery disease or diabetes, 2) presence of any clinically significant disease that may interfere with the results or the subject’s ability to participate in the study, 3) any major surgical procedure or trauma within four weeks of the first study visit, or 4) medication for hypertension or hyperlipidemia. During 2015-2016, the 101 subjects in SW3P visited the clinic every three months, four times in total. In 2016-2018, 97 subjects continued to visit the clinic two additional times with a gap of six months between appointments. Each visit included the measurement of body weight, waist and hip circumference, body fat using bioimpedance (Tanita MC-780MA), and blood pressure. Changes in health and life-style was recorded at each visit with a questionnaire covering factors such as diseases, infections, medication, exercise level, and personal perception of health. All 101 subjects were instructed to fast overnight (at least 8 hours) before the collection of blood, urine and feces. Human EDTA plasma samples were transferred on dry ice to SciLifeLab and stored at −80°C upon arrival. Clinical and demographic characteristics are presented in Table S5 and Table S6. Genome analysis is described in the Supplementary. The study was performed in accordance with the declaration of Helsinki and the study protocol was approved by the Ethical Review Board of Göteborg, Sweden (*Regionala etikprövnignsnämnden*, Gothenburg, Dnr 407-15, 2015-06-25).

### TwinGene samples

In the present study, serum samples from a set of 3,000 individuals from the TwinGene study [13] were used for validation purposes. The details about the study, the sample selection criteria and randomization, the plasma protein profiling and genome analysis of the samples are described in the Supplementary as well as by Hong et al [14]. The TwinGene study was approved by the Ethical Review Board (*Regionala Etikprövningsnämnden*, Stockholm, Dnr 2007/644-31).

### Sample preparation

Crude EDTA plasma was stored at −80°C. Prior to aliquotation, samples were transferred to −20° overnight, thawed at 4°C and then vortexed and centrifuged (3,000 rpm for 2 minutes). Using a liquid handling robot (EVO150, TECAN), samples were randomized across 96-well microtiter plates. Protein labeling was performed with biotin, as previously described by Drobin et al [11]. Briefly, EDTA plasma was diluted ∼1:10 in PBS and biotinylated with NHS-PEG4-Biotin (Pierce) dissolved in DMSO. Following 2 h incubation at 4°C, the reaction was quenched with Tris-HCl 0.5 M, pH 8.0. Prior to analysis, a small version of a protein profiling test was performed to confirm successful biotinylation. Labeled plasma was stored at −20°C until analysis.

### Antibody suspension bead array assays

We used a total of 1,450 antibodies raised against 896 unique protein targets, including 1,285 antibodies from the Human Protein Atlas (HPA) project [15], 72 mouse monoclonal antibodies (BioSystems International Kft) and 93 antibodies from other commercial vendor (Supplementary Data file S1). Each Suspension bead array (SBA) was assembled by covalently coupling antibodies to magnetic and color-coded MagPlex beads (Luminex Corp.) and mixing these beads to create the arrays. The procedures for antibody coupling and bead mixing can be found in Supplementary Methods and together with the following protocol for plasma profiling have been described by Drobin et al [11]. Briefly, 5 µl of the beads from one SBA was aliquoted into each of the wells of 384-well microtiter plates. Biotinylated plasma was diluted 1:50 in PVXC buffer, supplemented with 0.5 mg/ml purified rabbit IgG (Bethyl laboratories) and then heated at 56°C for 30 minutes. Forty-five microliters plasma was transferred to the bead plate and incubated with the beads overnight, during which time the antibodies bind their corresponding antigen in the sample. Low-affinity complexes and unbound proteins were removed in consecutive washing steps with PBS-T 0.05% (EL406 washer, BioTek). Beads were incubated for 10 minutes with 0.4% paraformaldehyde, washed, and then incubated for 20 minutes with R-phycoerythrin-labeled streptavidin 1:750 (Invitrogen). Lastly, the beads were washed and fluorescent signal from binding events were detected with a FlexMap 3D instrument (Luminex Corp.). Signal intensities reported as Median Fluorescent Intensity (MFI) were exported from the software xPONENT (Luminex Corp.) and at least 32 events per bead ID were used for data processing. As described in the Supplementary, a subset of antibodies was selected for statistical analyses based on performance in the SBA assay and orthogonal proteomics approaches including sandwich assays, mass spectrometry and proximity extension assays (Table S7 and Supplementary Data file S3).

### Statistical analysis

Data analysis and visualizations were performed using the statistical software R version 3.6.0 [16] as described below, with details provided in the Supplementary. All statistical and technical evaluations were performed using log-transformed MFI unless otherwise stated. To account for plate and batch effects, AbsPQN with Multi-MA normalization was applied by 96-well microtiter plate (Fig. S11). Spearman’s rho statistic was used for estimating the correlation between variables, unless otherwise specified. Manhattan plots were drawn by using qqman package (v 0.1.4). For the seasonal association analysis, a model for regular cyclic movements across time was fitted to each protein profile. UMAP analysis was performed on centered and scaled SBA data using the R package umap. As UMAP has several hyperparameters that can influence the resulting embedding, we compared if results were conserved with Euclidian distance while varying the set seed, n_neighbors and min_dist parameters, as exemplified in Fig. 4 with n_neighbors = 10 and min_dist = 0.25. Inter-class correlations were calculated with the R function ICC() from the psych package using ICC(3, 1) levels as output. Association between protein profiles and clinical traits was tested by linear-mixed models using the R package lmerTest [17] and visualized with the circlize package [18]. Protein profiles were standardized to z-scores and applying a linear model, three variables; intercept, slope and sum of residuals (absolute value) were calculated over time for each individual and protein. Protein modules were defined using the WGCNA v. 1.66 [19, 20] as described in further detail in the Supplementary.

## Results

### Study overview

We delineated the longitudinal characteristics of proteome profiles in a Wellness profiling cohort (denoted S3WP) of 101 individuals who donated plasma samples at four visits during one year (Fig. 1). With our antibody bead array data, we performed a series of data analyses on clinical, longitudinal, network and genetic aspects in order to capture the inter-individual diversity and longitudinal variability in the circulating proteomes. Further details about the experimental design can be found as Supplementary Information.

**Fig. 1.**
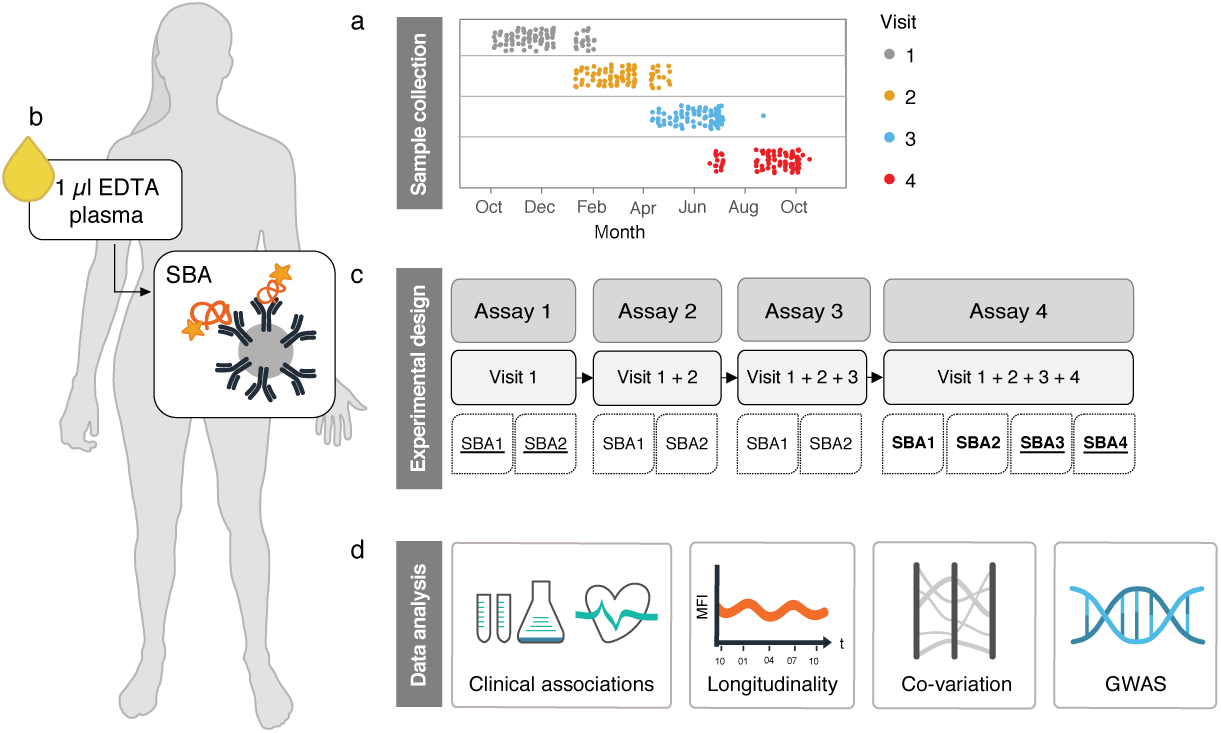
Experimental design and data analysis pipeline. (A) Over the course of one year, samples from 101 individuals were collected at four different visits to the clinic. (B) Plasma proteins were measured from 1 µl EDTA plasma with antibodies conjugated to beads. (C) Following each completed visit, samples were randomized within an assay and analyzed together with all previously collected visits. In total, four SBAs were created and incubated with the samples as indicted in the flowchart. (D) Protein profiles were tested for associations to clinical traits, longitudinal stability, networks of co-regulation and GWAS. The underlined labels correspond to assays where the complete set of samples were analyzed in duplicate. Labels in bold correspond to assays where the SBA was incubated with 96 replicated samples for technical validation. SBA, suspension bead array; GWAS, genome wide association study.

### Annotation of antibody-derived protein profiles

First, we selected the most reliable antibodies from the initial set of 1,450 antibodies targeting nearly 900 unique proteins (Supplementary Data file S1, Fig. S1). As described in the Supplementary in further detail, applying a combination of validation criteria led to the selection of 734 protein profiles (Table S1, Table S2, and Table S3). This assessment included the use of genome wide association studies (GWASs) to identify single nucleotide polymorphisms (SNPs) in the protein-encoding regions of the target genes (Table 1 and Table S4). In summary, we annotated the 1,450 antibodies included in the assays and selected 734 unique protein features for further investigations.

**Table 1.**
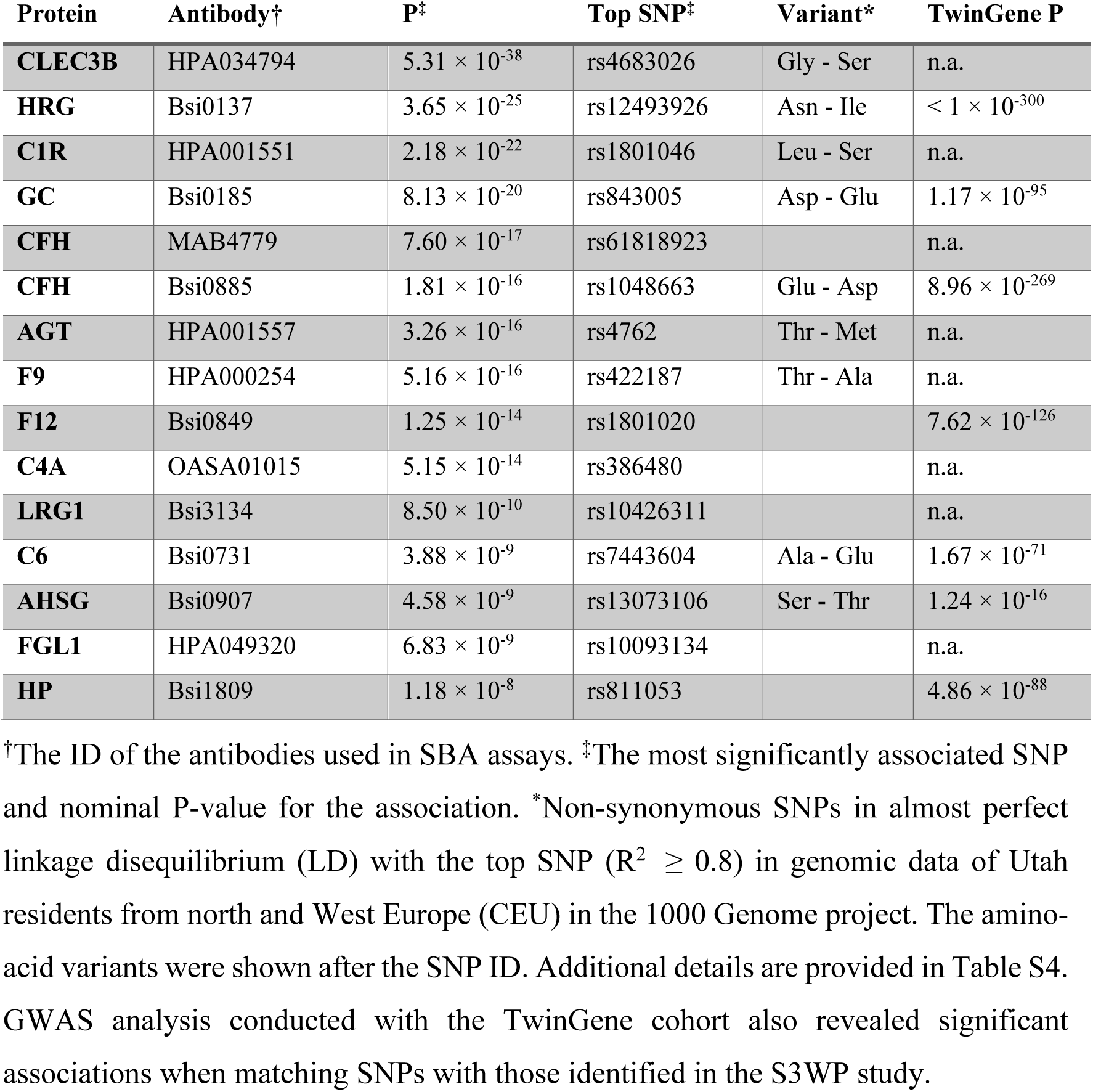
Proteins with *cis*-pQTLs.

### Clinical associations of circulating proteins

First, we referenced the 734 protein profiles to the clinical traits measured with standardized clinical tests, BMI and smoking habits. Applying linear-mixed effect models, we identified statistically significant protein-trait associations (FDR P ≤ 0·001). As shown in Fig. 2 and Fig. S2, these associations were enriched for traits like triglycerides (TG) (n = 42), CRP (n = 32), apolipoprotein B (ApoB) (n = 21), total cholesterol (Chol) (n = 19), low-density lipoprotein (LDL) (n = 28), and the ratio of ApoB/apolipoprotein A1 (ApoB/ApoA1) (n = 13). As expected, strong associations were seen between clinical and proteomic CRP (FDR P = 3·47 × 10^−160^) and ApoB (FDR P = 3·90 × 10^−25^). We discovered other strong associations that included TNFRSF1B and DAPK1 with TG (FDR P = 3·01 × 10^−55^ and P = 5·21 × 10^−48^). Other top associations for the clinical traits (FDR P < 1 × 10^−3^) were for BLVRB to hematocrit (Hct); THBS1 to platelet count (Plt); S100A9 to the count of white blood cells (WBC) and neutrophils (Neut); SAA and FGL1 to CRP; LEP to BMI; ANGPTL3 to ApoA1; RARRES2 to cystatin C (CystC); CCL16 to of gamma-glutamyl transferase (GGT); and IGFBP2 to levels of N-terminal pro B-type natriuretic peptide (NTproBNP). This showed that a variety of expected associations were replicated in our proteomics approach, many related to secreted proteins, and that associations to the traits related to inflammation and lipid metabolism were most prominent.

**Fig. 2.**
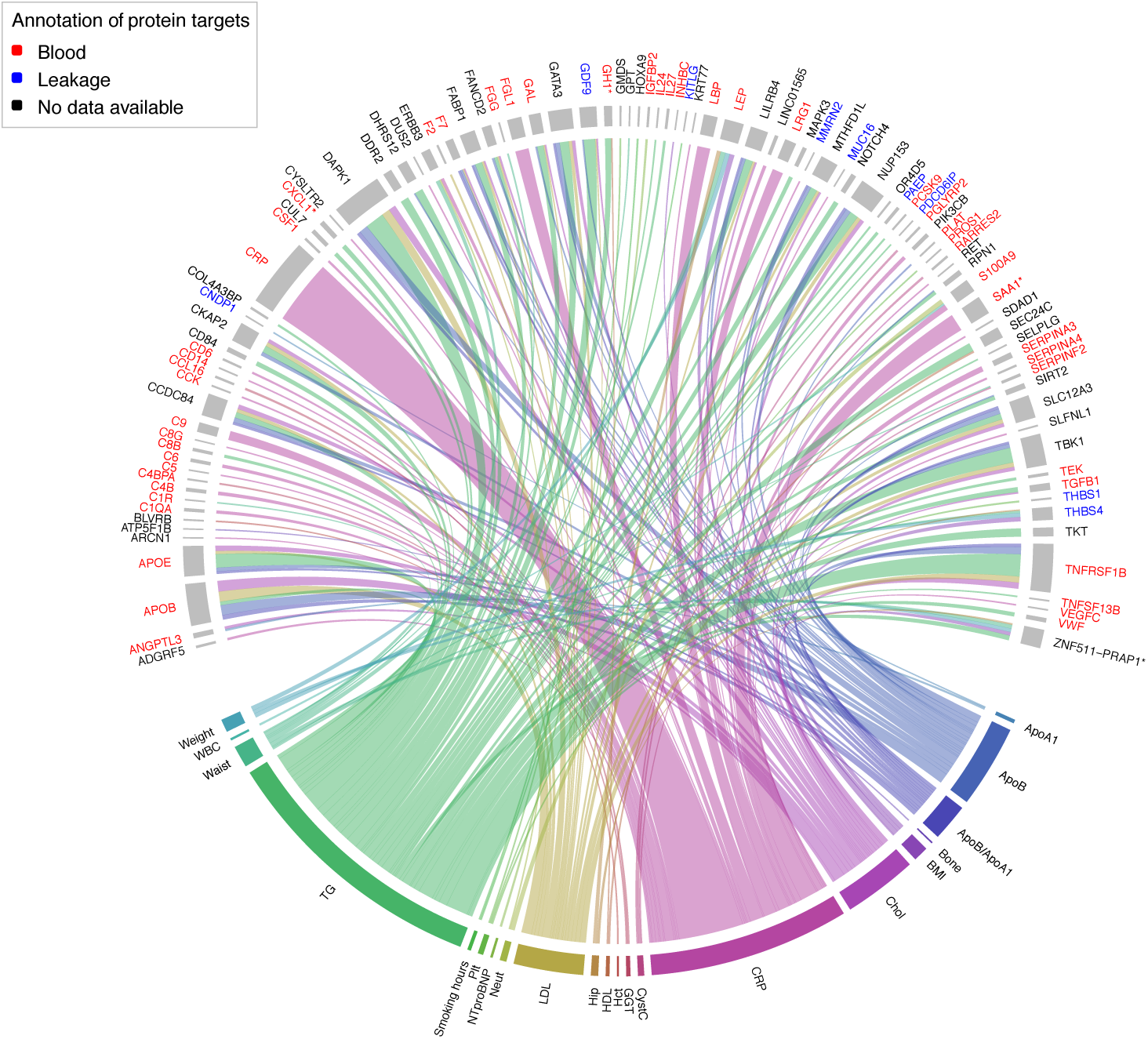
Association map of proteomics and clinical traits. Chord diagram of associations (FDR P < 0·001) between protein profiles and clinical traits obtained from linear mixed effect models. Line thickness is proportional to -log10(P-value) and colored by clinical trait. Protein features that represent a family of several proteins are denoted with one gene name followed by “*”. Feature names are colored red if predicted to be actively secreted into blood, or blue if they appear in blood due to cell leakage [9, 39].

### Assessment of longitudinal protein profiles

To capture longitudinal changes across the four consecutive visits, we investigated the reproducibility of protein measurements across repeated assays (Fig. 1). We performed inter-assay correlations of the protein levels (“technical variability”) and compared these to inter-visit correlations (“longitudinal variability”). Intraclass correlations (ICC) were computed for both measures and allowed us to consider the technical variability when judging the longitudinal variability. Protein profiles with ICC ≥ 0.8 were defined as technically consistent and/or longitudinally stable. Out of all protein targets, 61% (447/734) revealed a high technical stability and 58% (428/734) were stable longitudinally. A total of 49% (359/734) of all proteins could be measured with a high precision when including both the inter-visit and inter-assay ICC ≥ 0.8. Reassuringly, the distribution of ICCs obtained from the proteomics approach was similar to the values obtained from the clinical tests (Fig. S4A, Table S5, Table S6).

The most consistently measured and longitudinally least variable protein was CD5 molecule like (CD5L, inter-visit ICC = 0.97) as exemplified by its technical and longitudinal profiles shown in Fig. 3A and Fig. S4B. On the other end, Caldesmon 1 (CALD1) was one of the proteins with high technical precision (inter-assay ICC = 0.88) but also a high variation between consecutively collected samples (inter-visit ICC = 0.32), as shown in Fig. 3B and Fig. S4C. This aligns with our previous findings of CALD1 being susceptible to conditions related to plasma preparation [21]. In summary, we found that ∼50% of the proteins were measured with high precision and low longitudinal variability in blood plasma throughout one year. An additional analysis of seasonal effects on the plasma proteome, see Fig. S5A-C and Supplementary for details, only found levels of FLNA and BLVRB to fluctuate with season (FDR P < 0·01).

**Fig. 3.**
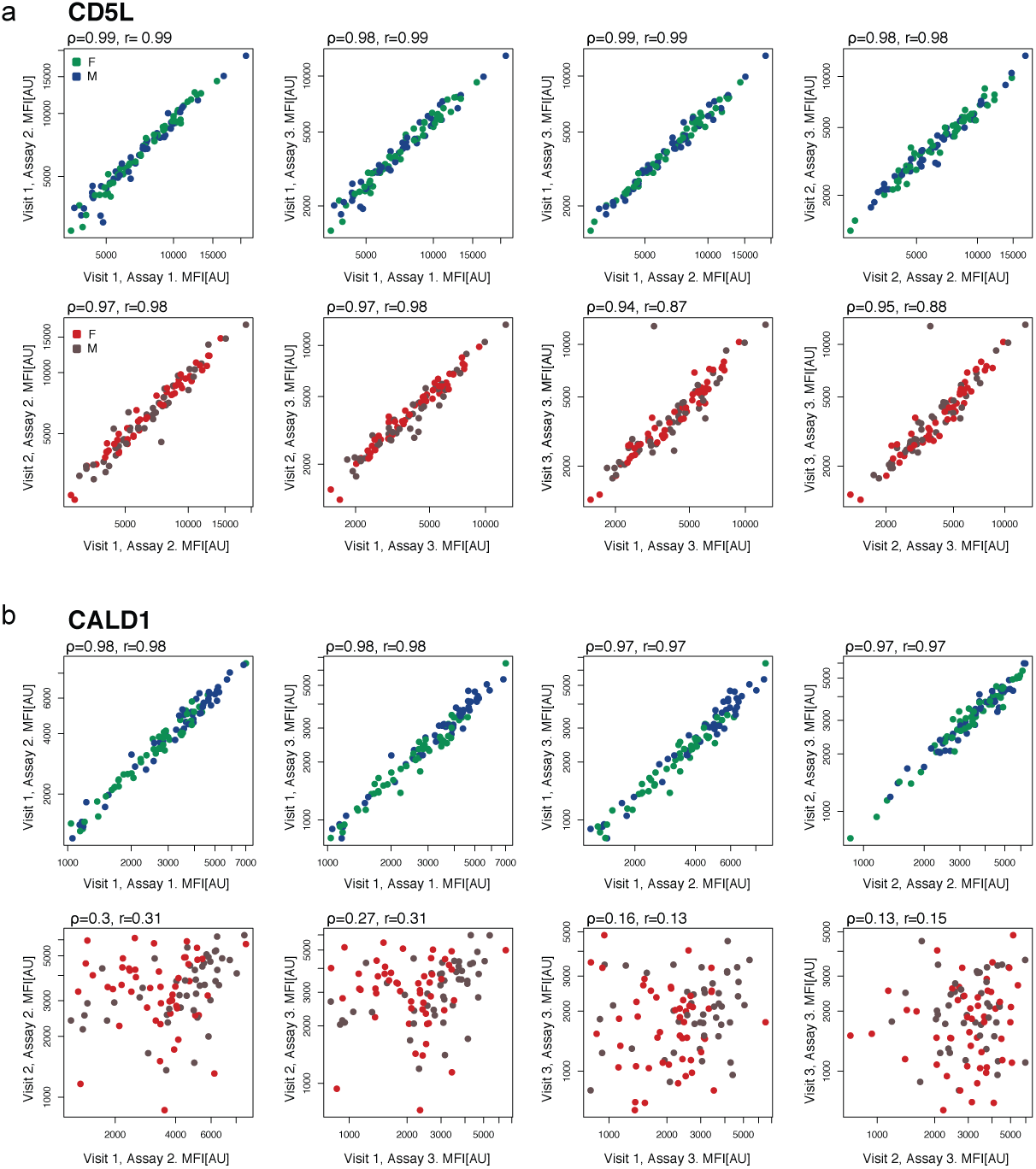
Inter-assay and inter-visit variability. Shown are correlations of technical (inter-assay, upper panel) and longitudinal (inter-visit, lower panel) profiles. (A) CD5 molecule like (CD5L) represents a both technically and longitudinally stable protein, while levels of (B) Caldesmon 1 (CALD1) vary between visit but not repeated assays. Each dot represents one individual, colored by sex (F, female; M, male), MFI relates to median fluorescent intensity and AU are arbitrary units.

### Global analysis of protein profiles

Next, we investigated if the combination of protein profiles contributed to personal plasma proteomes signatures. We used Uniform Manifold Approximation and Projection (UMAP) [22] to compress the data from all 101 samples and 734 proteins into two dimensions (Fig. 4). Each subject clustered predominantly with itself across all four visits. This implied that the plasma proteome signatures were diverse and composed of unique combinations of protein profiles for each individual participant. Computing the individual longitudinal variability per participants revealed ICCs = 0.99 ± 0.005 (mean ± SD). This highlights the existence of a stable and person-specific proteome signature, but also suggests that there is a considerable diversity in the circulating proteomes between clinically healthy subjects.

**Fig. 4.**
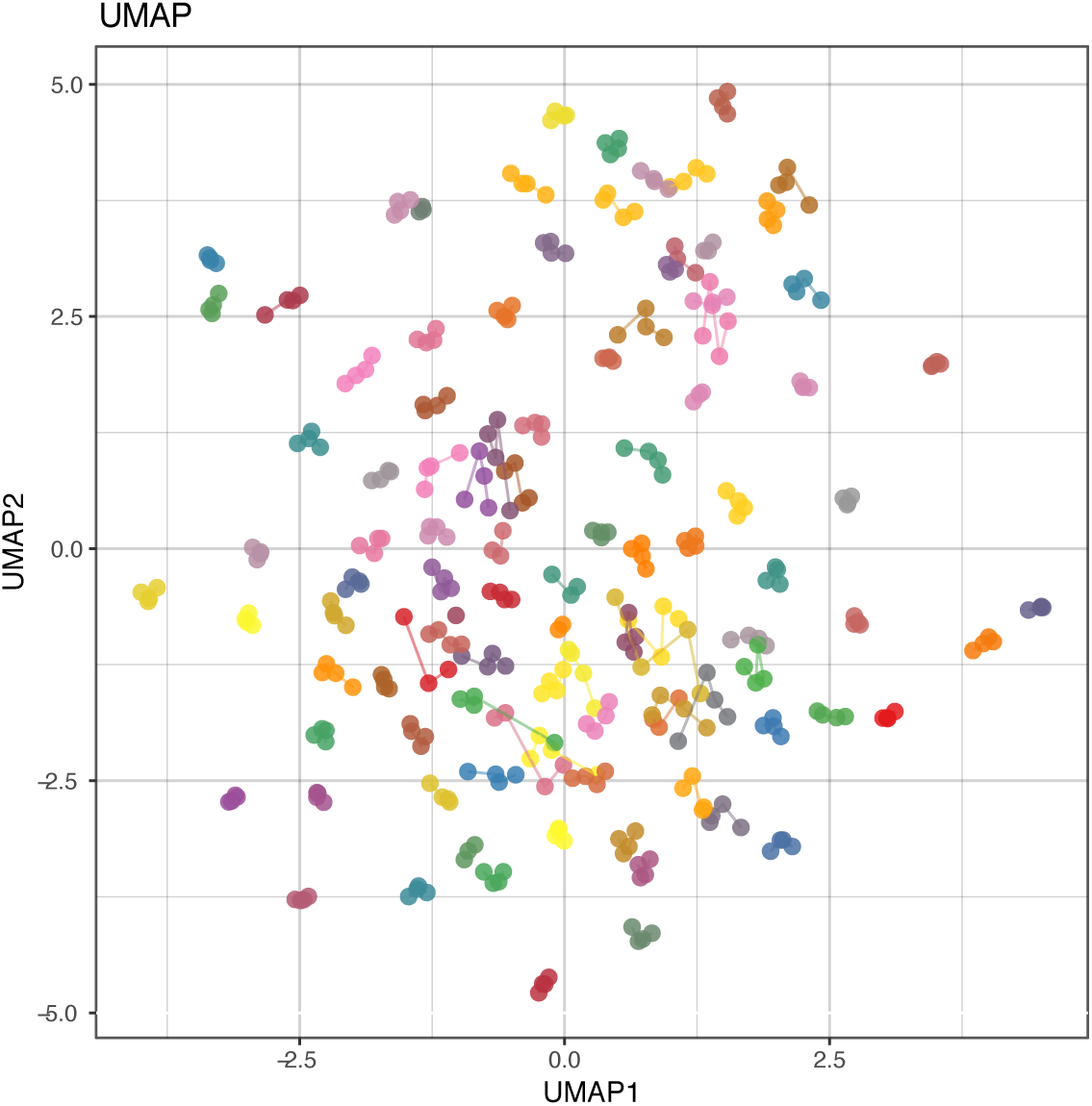
Diversity of individual-specific protein profiles. UMAP analysis of 734 protein features and samples from four visits, colored by subject (N = 101). Colored lines indicate which samples belong to the same individual. UMAP, Uniform Manifold Approximation and Projection.

### Longitudinal co-regulation of plasma proteins

In addition to investigating each protein individually, we explored if there were longitudinal networks of co-varying protein profiles. We used weighted gene co-expression network analysis (WGCNA) to define, annotate and analyze modules of co-regulated and interconnected protein profiles (see Supplementary material for details).

Computed for each visit, WGCNA resulted in eight mega modules (Fig. S6) and each mega module contained 11-242 proteins. We then tracked the mega module membership across all visits to create a map of the longitudinally patterns of conserved “core modules” (Fig. S7). From the eight mega modules per visit, we created core modules by matching all possible combinations of sequential overlaps between the mega modules across the visit (Fig. S8). A protein was ultimately assigned to one of the core modules if it was part of a particular pattern across all visits. Eight out of nine core modules contained at least one protein, and 59% (434/734) of the proteins could be assigned to one of these eight core modules (Fig. 5).

**Fig. 5.**
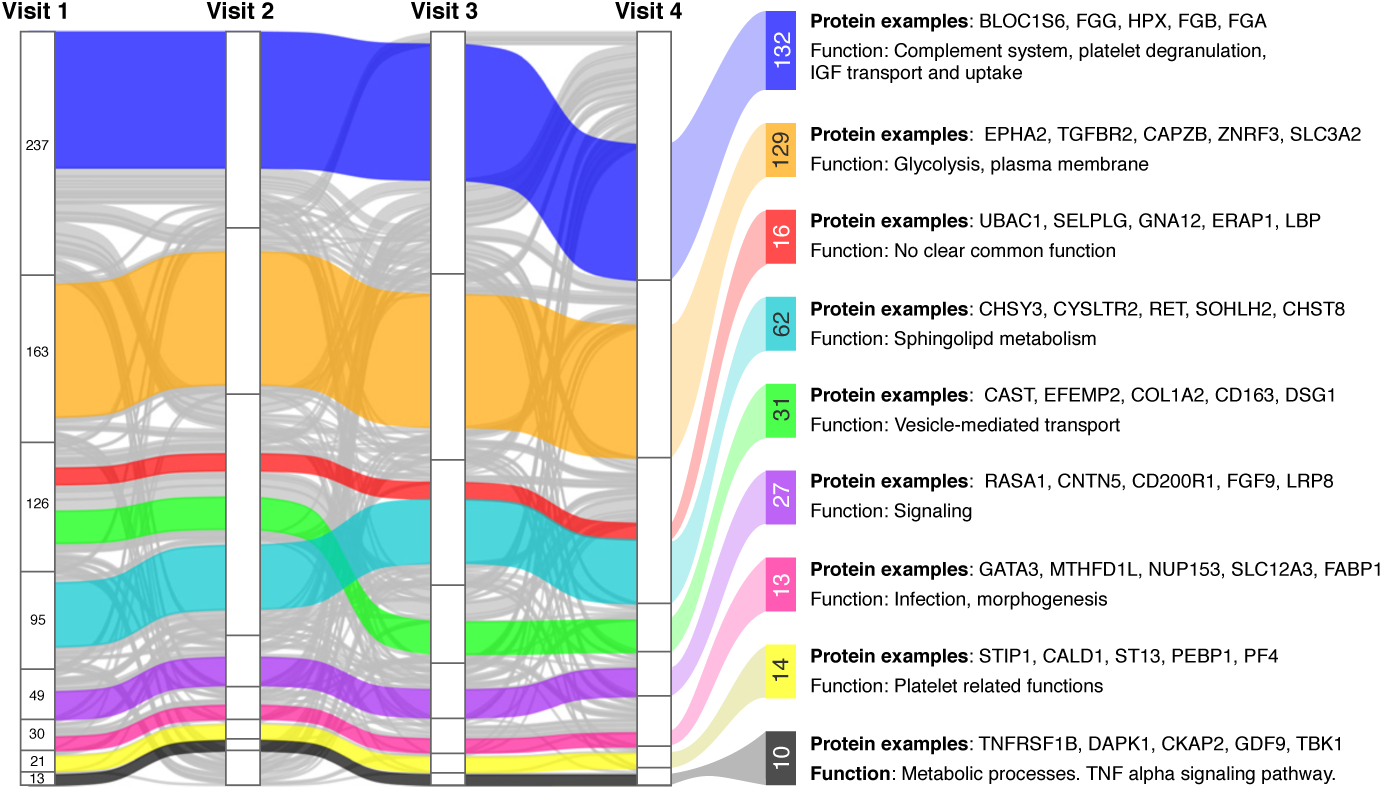
Networks of co-varying proteins. WGCNA was used to determine co-varying proteins per visit (stacked groups) and across visits (horizontal bands). Each vertical line represents one protein and its mega module membership in each visit. Proteins are colored according to the core pattern they belong to. Proteins that do not belong to any core pattern are grey. Each core pattern is annotated to the right with the number of proteins it contains, a summary of associated pathways and GO terms, and examples of proteins following the given pattern. The examples of proteins given are the five proteins with the highest correlation to the core pattern eigengene.

The eight core modules were then annotated for their biological functions and their relation to clinical traits, as it is expected that proteins within the same core module could share biological functions and interactions, or can be controlled by common mechanisms (Fig. 5, Supplementary Data file S2). This revealed associated pathways and annotations related to biological functions like complement system, vesicle transportation, platelets and metabolic processes. Next, we explored links between groups of co-varying proteins with the available clinical traits. We identified a number of statistically significant associations (FDR P < 0·01) for the blue and black WGCNA core modules that correlated with lipid related traits, but in opposite directions. The blue pattern was negatively correlated with the levels of triglycerides and the fraction of ApoB and ApoA1, and conversely the black pattern was positively correlated with these traits as well as levels of ApoB and LDL. Additionally, the blue pattern was negatively correlated with CRP. Thus, these two different sets of co-regulated proteins likely have opposite functions within lipid metabolism by being linked to LDL and HDL respectively. This is consistent with the fact that the LDL associated protein ApoB follows the black core pattern and the HDL associated proteins ApoA1 and ApoA4 follow the blue pattern. None of the other core modules had significant associations to the available clinical traits, even though the effects were mostly consistent across visits. None of the core modules had significant associations to sex or age.

In summary, we found longitudinally conserved modules of protein co-expression networks with associations to biological functions and clinical traits.

### Genetic effects on the plasma proteome

As introduced above, we used genetic data and found 15 *cis*-protein quantitative trait loci (pQTLs) for 14 protein profiles (P < 1·35×10^−8^, Bonferroni P < 0·05) from the association tests with non-redundant ∼3.7M SNPs (see Table 1, Table S4 and Fig. S3). All 14 unique proteins were annotated to be secreted into blood and primarily expressed by the liver [9] and the longitudinal profiles stratified by genotypes are shown in Fig. 6A-O. Following our previous observations [14] and recent insights connecting the circulating proteome with genetic variation [23], we investigated if differences in detected proteins can be linked to protein polymorphisms. Even though non-synonymous SNPs are rare (< 0.3%) [24], we found these to be overrepresented among the identified *cis-*pQTLs. Indeed, out of the 15 identified pQTLs, nine of the loci (60%) contained variations that induce a change of amino acids in the protein sequence. Because these genetic associations were obtained in the relatively small S3WP cohort of 101 subjects, we corroborated the results by checking for the same associations between genetic variation and circulating proteins in an independent set of 3,000 individuals from the TwinGene study [13].

**Fig. 6.**
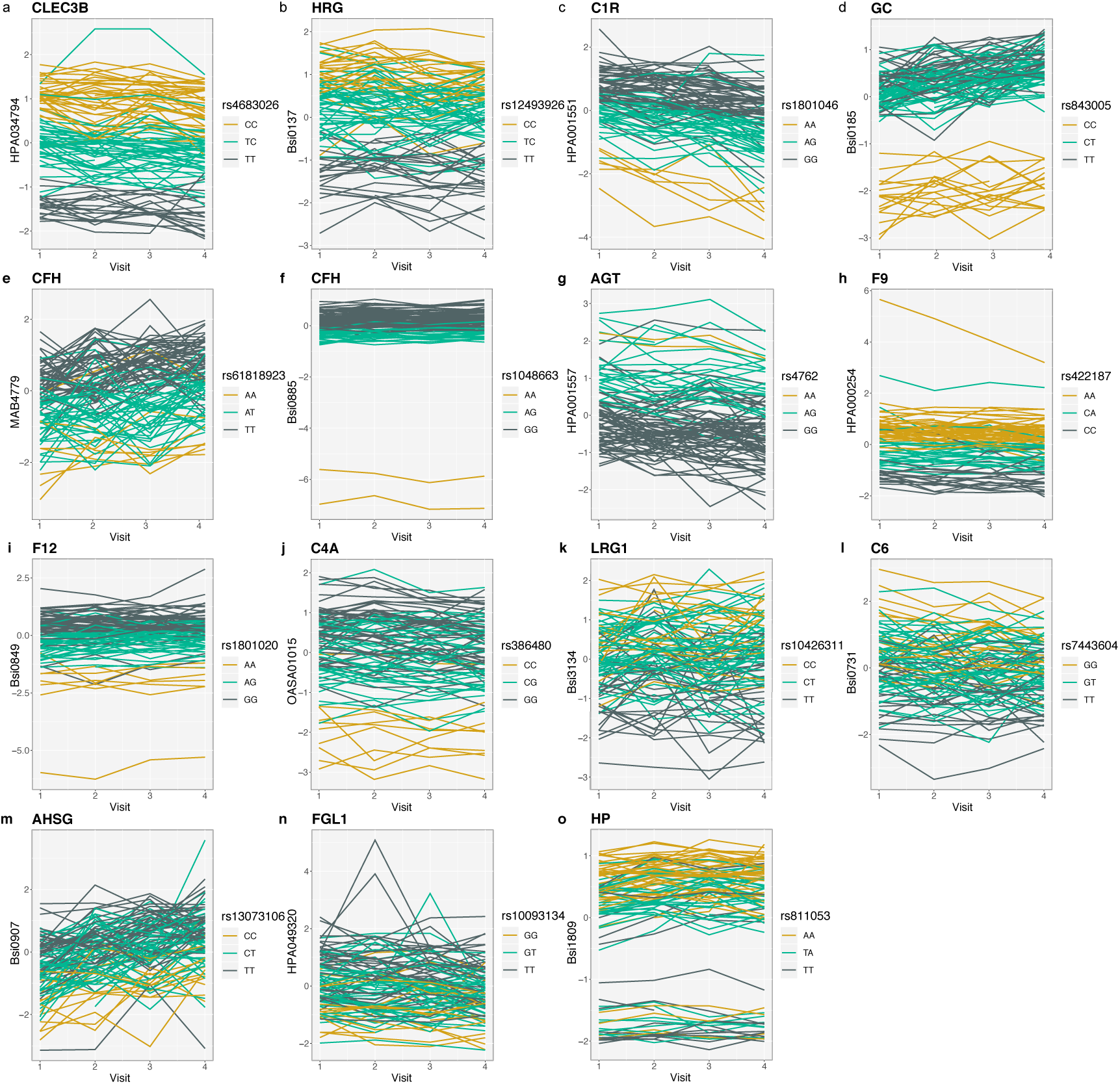
Longitudinal characteristics of plasma protein genotypes. The line plots show plasma proteins associated to genetic variants where z-scores were used to represent protein levels. Each line represented one individual and color codes the genotypes. Only individuals with data from all four visits were included for visualization.

Among the most significant association between genetic variation and circulating proteins were Tetranectin (C-type lectin domain family 3 member B, CLEC3B) and the SNP rs4683026 (P = 5·31 × 10^−38^). One of the perfect linkage disequilibrium (LD) proxys (R^2^ = 1) of that SNP is rs13963, which can lead to two proteoforms with either a serine at the 106^th^ position or a glycine. Detected levels of CLEC3B, a protein secreted by the lung, muscle, spleen, and adipose tissue, were highest for the CC genotype and decreased as the number of C alleles decreased (Fig. 6A). This indicated that the assay preferred the Gly106 isoform produced by the C allele over the Ser106 isoform produced by the T allele.

Similarly, proteoforms of liver secreted vitamin-D-binding protein (GC) could be linked to non-synonymous SNPs, hence reporting an isoform-specific affinity rather than differences in abundance [25]. We indeed found that specific alleles had major effects on the reported amount of this circulating protein (Fig. 6D). We concluded that the levels of GC detected by the assays were strongly determined by the genetic variants rs222047, rs843005 and rs7041. Reassuringly, rs7041 had previously been described as a *cis*-pQTL of GC when using an even larger study set and another type of quantitative immunoassay [26].

As shown in Fig. 6I, we also found a *cis*-pQTL SNP rs1801020 (P = 1·25×10^−14^) corresponding to the 5’ untranslated region of the coagulation factor XII (F12) gene [27], and we replicated this association in the TwinGene cohort (P = 7·61×10^−126^). The common genetic variant rs1801020 modulates F12 liver expression [28], and thus provides additional evidence that the detected SNP modulates gene expression, which in turn impacts the F12 protein abundance rather than the protein sequence. Out of all 101 subjects, we found that among seven participants with the F12 genotype, one individual had substantially lower secreted levels of F12. Subsequent tests of the participant in the clinic confirmed a delayed activated partial thromboplastin time (aPTT), which has also previously been described for this F12 polymorphism [28].

Besides GC and F12, we identified subgroups of individuals linked to differences in plasma protein levels for the secreted liver proteins complement factor H (CFH) and haptoglobin (HP). Lower levels of HP were determined in 22% (21/93) of the plasma samples from the S3WP study participants and in 15% (447/2,974) of the sera from TwinGene study (Fig. S9A). For one of the anti-CFH antibodies (Bsi0885), the detected protein levels were lower in plasma of 2% (2/93) of the S3WP individuals, and equally in 2% (62/2974) of the TwinGene participants (Fig. S9B). Compared to the second anti-CFH antibody (MAB4779), which was not included when analyzing the TwinGene samples, the main SNP for Bsi0885 did induce a missense mutation affecting the protein’s sequence. This effect could explain the differences in binding properties of the antibodies towards the variants of CFH. No distinct population subgroups with either lower GC or F12 protein levels were detected in the serum samples of the TwinGene study (Fig. S9C-D). As further described in Supplementary, we also compared pQTLs with eQTLs and other RNA expression data to pinpoint the source of expression regulation of proteins with *cis-*pQTLs. There were no significant associations between the genetic data and clinical.

In summary, distinct differences in plasma protein levels can be explained by genetic variants. These insights are valuable when comparing the protein levels between individuals as they can provide another motivation for why a more precise and personalized assessment of health in circulation requires both longitudinal monitoring and the influence of genetics.

### Facets of individual and longitudinal protein profiles

UMAP analysis revealed that overall, person-specific profiles remained stable over time. To identify inter-individual differences on a protein level, we z-scored the data and determined the inter-quartile range (IQR) as a measure of diversity between individuals. Only minor differences between the participants of the study where seen for proteins such as DSC3, GFAP, and GDF15 (IQRs ≤ 0.15), as compared to more prominent inter-individual diversity in levels for the liver proteins LEPR, IGFBP2, FCN2 or SERPINA1 (IQRs ≥ 1.5).

To further illustrate changes occurring in the plasma proteomes, we queried the data for representative examples among the participant’s protein profiles (Fig. 7). We asked which protein might vary due to distinct events, remain different during the study, or gradually change over time in any of the individuals. We again used protein z-scores with all participants serving as a reference population. To this end, we selected the 359 most stable protein profiles (ICC ≥ 0.8) in order to focus on capturing individual rather than common patterns. We applied an annotation scheme that was based on the parameters we obtained from fitting linear models to every protein profile. We scored each of the protein profiles for every individual based on three criteria:

**Fig. 7.**
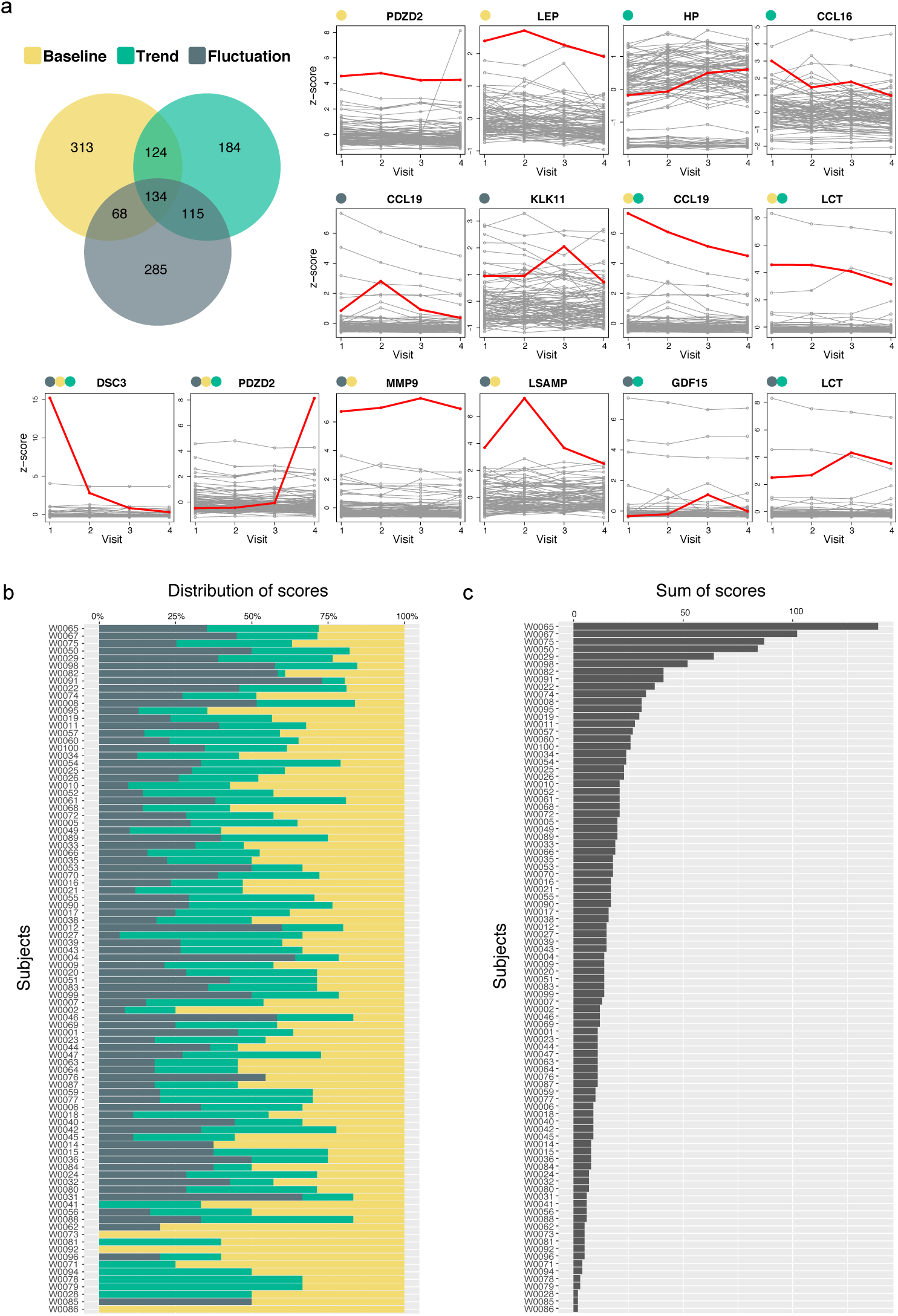
Facets of longitudinal protein variability. Protein profiles were stratified by their longitudinal profiles. (A) Venn Diagram indicates the number of observations (protein per individual) that was deviating (± 3xSD) from the population mean in terms of protein baseline, trend and fluctuations. Here, we selected 14 protein profile examples. Each gray line represents one individual. One selected individual with a particular protein profile is highlighted in red, and the category of the red profile is marked on the left side of the protein name. (B) Distribution of the three annotation criteria per subject, and (C) the sum of the three annotation criteria per individual.

i. baseline = individual deviation of protein levels from the population,
ii. trend = a person’s changes in (increasing or decreasing) protein levels,
iii. fluctuation = fluctuation of protein levels as deviation from linear changes.

In total, 33,028 profiles were derived from 359 proteins and 92 individuals. We classified each profile to each criterion as deviating if the obtained values were ± 3xSD of the population average. As shown in Fig 7, 3,7% (1,223/33,028) of all possible participant-protein measurements revealed a variation at the individual level on one or several of the categories. This frequency is ten times higher than observing these variations by chance (0,27%). We then summarized these scores and evaluated the outcome per protein (including all individuals) and per participants (including all profiles).

Concerning the proteins, levels of CAPZB, RPSA, FGFR1, BPGM, and PECAM1 were longitudinally preserved and least unique to any participant as these were neither elevated, changed nor fluctuated over time. Interestingly, none of these proteins were annotated to be secreted into bloodstream, hence likely represent leakage products. In contrast, we found that secreted blood proteins GDF15 and MMP9, as well as LCT, were variable proteins along the longitudinal axis when considering the total number of individuals with changes to any of the three annotation criteria. As exemplified by GDF15, aspects of inter-individual diversity and longitudinal variability can present as independent characteristics of circulating proteins.

Among the protein profiles of each participant, there was at least one protein with elevated baseline or trends of fluctuating levels (Fig. 7B-C). The most variable profiles were found for participant W0065, as there were 39 proteins being different in term of baseline levels, 51 increased or decreased, and 49 proteins fluctuated over time. In contrast, participant W0086 was ranked as most stable (accumulated score = 2) and the two deviating profiles corresponded to elevated baseline for the two proteins PDZD2 and KLK11.

This analysis illustrated that there is diversity in terms of how longitudinally stable or variable an individual’s plasma proteome can be. It is likely that many of the unique and stable trajectories that deviate from the average population baseline might be due to genetic effects, lifestyle factors or medication, however our study was underpowered to extract other associations of weaker effect sizes.

## Discussion

We profiled 101 individuals using a multiplexed affinity proteomic assay and found that the plasma proteome signatures were highly individual-specific. To address concerns about antibody validation, our exploratory multiplexed approach applied a scoring scheme to identify binders with consistent performance across assays and longitudinally collected samples. We highlighted findings related to individual protein variability, found interesting links to genetic components and networks of co-regulatory proteins, and lastly demonstrated the potential benefits of individual-level, longitudinal protein profiling. Our observations can have substantial impact on studies searching for common disease proteins across a population, because both the inter-individual diversity and longitudinal variability can have influence on the composition of the circulating proteome.

Affinity binders are important tools frequently used in research and as diagnostic reagents. The current concern surrounding the reproducibility of data derived from such research has raised awareness and increased the efforts in developing strategies for antibody validation [29]. The utility of antibodies is context dependent and the performance may vary depending on technological method or sample composition. Therefore, it is necessary to annotate and validate the specificity of each binder in its intended application, preferably by using orthogonal methods as applied here. The selected antibodies have passed several validation steps in the generation pipeline [30], but validation needs to be tailored for plasma analysis [31, 32]. The multiplexed assay applied here was an exploratory effort allowing the analysis of large numbers of samples and analytes. The method relied on a single binding event between an antibody and its target protein, hence, the inherent risk of the method is off-target binding, unspecific binding, or capturing of protein complexes [33]. Notably, several proteins measured with our assays were identified with a *cis*-pQTL, providing inferred evidence for on-target binding. By combining information for each antibody regarding technical reproducibility and supportive data including other antibody-free assays [34], we developed a transparent annotation strategy to limit our analysis from the initial 1,450 antibodies to a set of ∼700 high confidence protein profiles of which almost 50% were very stable over time. Indeed, many proteins in the latter selection can be expected to be detectable in the circulation because these were either secreted from solid tissues into blood or leakage from blood cells.

Considering the intra-individual diversity, we found that ∼50% of the studied protein profiles were stable across the four sampling occasions over one calendar year. We acknowledge though that some proteins can appear to be more dynamic depending on the studied timespan, or if any other perturbation, disease or intervention occur among the participants. For the majority of the proteins, variability was low for both the technical replicates and between time points. Hence, the fluctuations observed between two time points could also likely be due to technical variability rather than life style related perturbations. Nonetheless, changes that occurred on a continuous scale for a subset of the study group could serve as supportive evidence for physiological rather than technical changes. Clearly, our observations are restricted to the proteins targeted by our assays and may be affected by limitations in terms of the sensitivity to the effects from protein interactions as well as co-enrichment of other proteins [33]. Nonetheless, we observed a high consistency when profiling circulating proteins in the longitudinal sample collection, which encompasses the process of drawing blood, processing the collections and analyzing different samples from the same subject. Hence, a longitudinal assessment reflected a combination of different variables such as sampling, biobanking, assays and physiological changes. However, we found and focused on protein profiles with low longitudinal variability. This indicated that the applied concept, sampling schemes and method had the precision to define personal baseline values and capture individual changes.

We applied multivariate analysis to cluster 734 protein features from four time points and found individual-specific profiles that were retained throughout one year. This observation was further supported by the fact that the majority of proteomic and clinical profiles showed ICC > 0.9 between the visits for each individual. It is worth noting that our study included a small number of subjects (∼100) followed during a relativity short time span (one year). Further, the participants were deemed clinically healthy with a balanced (and possibly more deliberately healthy) lifestyle during this period. Both the intra-individual diversity and the longitudinal variability are important observations because these can lay the foundation to a next-generation of studies aiming to more accurately assess diseased individuals or those in treatment. It will, however, require even larger study populations that include a defined intervention, such as common disease incidents, drug treatment, or surgery to determine which subset of proteins are needed for a respective disease phenotype.

We also applied a network approach to study any coordinated change of several proteins over time. Although proteins within the same WGCNA-defined module were covarying, we did not collect evidence about their physical interactions. These modules rather suggest that there is an interconnection that can coordinate protein expression. Hence, and as previously observed by others on single time point measurements [35], there are possible processes that co-regulate protein levels via common mechanisms. Indeed, most of the identified core patterns were significantly enriched by proteins related to a particular biological function such as lipid metabolism (LDL, HDL and triglycerides). An added value of longitudinal profiling was further illustrated by WGCNA because not all proteins that co-varied within a single time point also continued to do so across all visits. Our findings suggested that coordination of other disease related networks and processes exists, but these may require a more dedicated study design and include pre-selected proteins. Nonetheless, we demonstrated in a study of clinically healthy individuals that processes related to metabolism, coagulation and inflammation were among the major coordinated functions of the plasma proteome and that these should be considered in any assessment of human health states.

Profiling the plasma proteomes identified groups of participants that presented with distinct differences in circulating protein levels. These subgroups could be linked to *cis*-pQTLs such as for the proteins GC, F12, CFH and HP. Investigating the identified SNPs, we found that the variation at the loci for GC and CFH coded for a missense mutation. This implied that the assay measured the relative abundance of specific proteoforms rather than detecting different concentration levels. We explained this by changes inducted to the sequence, structure or even post-translational modifications that will make the antibodies bind to each of the proteoforms with a different affinity. Reassuringly, we observed concordant associations even in serum samples of the TwinGene cohort, which we used as a validation set and that consisted mainly of elderly individuals.

One striking observation from a precision profiling perspective was to find a single participant with deficiency in F12 among all 101 individuals, and being able to use proteomic and genetic information to pinpoint a possible mechanism of lower levels of circulating F12. A deficiency in F12 is rare and generally non-symptomatic, however, in vitro F12 deficiency results in prolonged activated partial thromboplastin time (aPTT) [36]. Since aPPT is a measure of this and is a common screening test for hemostatic function, an underlying unknown F12 deficiency can have clinical consequences for the patient through inhibited or delayed invasive procedures or surgery and extensive diagnostic workup before a clinically relevant hemostatic disorder is excluded. Furthermore, common variants of F12, not resulting in deficiency, have been correlated with aPTT, presumably through modulating F12 levels [37]. A patient would therefore benefit from knowing about such a deficiency prior to surgery. The case of F12 illustrates how our proteomics data from continuous monitoring of a particular parameter can be combined with genetic data to generate information with direct clinical utility.

Collecting the pQTLs also allowed us to annotate whether differences in protein profiles were due to missense variants in protein coding regions or rather affecting gene expression. We used eQTL data accessible on the GTEx portal [38], accepting that the data is derived from tissues and cells from other individuals than the ones included in this study. Ideally, transcriptomic data from the same individuals should be incorporated in future analysis. Nonetheless, we found eQTLs in the liver (LRG1; F12), the artery (C6), pancreas (FGL1) or thyroid (C4A). Similarly, the relation of the pQTLs and splicing QTLs (sQTLs) were studied to annotate circulating proteins levels in relation to alternative splicing. We found sQTLs in the liver (AHSG; CFH), adipose tissue (CLEB3B), spleen (CFH), and thyroid (C4A). Connecting information from the pQTLs provides a useful approach to further annotate the levels of circulating proteins, even in 101 individuals. Nonetheless, the ∼10 proteins we have discussed point at the value gained when connecting proteomic with genetic data, such as for defining patient-specific cut-offs for disease classifications. This awareness will further assist our understanding of assay specific data in the context of precision phenotyping.

Lastly, we investigated the longitudinal variability of each protein across time at the level of each individual. We dissected this into three distinct categories of longitudinal variation that occurred among the participants. First, we identified individuals with increased or decreased baseline abundance of proteins that are consistent throughout one year. It can be hypothesized that baseline levels above or below a relative population mean might be due to genotype, and that other factors such as medication can further influence these. Next, we found proteins with gradually increasing or decreasing abundance across time. Monitoring the progressive changes across the consecutive sampling can highlight which proteins play a role in pre-symptomatic manifestation of a condition, or they reveal how effective a treatment of chronic conditions has been. Lastly, we found proteins that increased or decreased during shorter terms as these were captured only during specific visits. It remained a challenge to link many of these perturbations to reported changes in health, life style or behavior of an individual. Here a more detailed integration of the data on an individual level as well as decomposing the aspects related to sampling, shipment and analysis might be required. However, this demonstrated the importance to follow plasma proteomes over time in order to assess where on the spectrum of inter-individual diversity and longitudinal variability a specific individual resides. The distinction between time-resolved events and consistently changing or deviating baselines will consequently be important aspects to consider when implementing blood-based protein measurement to assess health in a clinic setting. It is by multiple layers of interconnected data, longitudinal sampling and individual-specific assessment over time that we can start predicting protein trajectories in time. Hence, utilizing such collected information will enable to distinguish between life-style related and short-lasting events (e.g. stress) over physiological processes that point at the onset, progression or manifestation of a disease or condition.

In conclusion, we profiled longitudinal plasma samples from 101 subjects using exploratory affinity assays and found that proteome profiles of clinically healthy individuals were diverse and highly individual-specific. While there were proteins varying over time in some individuals, many of the circulating proteins as well as their co-regulated networks were predominantly stable in this study population. Our work highlights the facets of individual-specific proteomes and the need to consider both inter-individual diversity and longitudinal variability when assessing health or disease states.

## Funding sources

This work was primarily supported by the Erling Persson Foundation for the KTH Centre for Precision Medicine (MU), and the Swedish Heart and Lung Foundation (GB). We also acknowledge the Knut and Alice Wallenberg Foundation for funding the Human Protein Atlas project (MU), and Science for Life Laboratory for Plasma Profiling Facility (JS). The Swedish Twin Registry is managed by Karolinska Institutet and receives funding through the Swedish Research Council under the grant no 2017-00641.

## Data and material availability

The S3WP datasets used for this report have been deposited with the Swedish National Data Service (www.snd.gu.se, a data repository certified by Core Trust Seal). The dataset can be made available for validation purposes by contacting. Data access will be evaluated according to Swedish legislation. Data access for research related questions in the S3WP program can be made available by contacting the corresponding author. Researchers interested in using Swedish Twin Registry data must obtain approval from a Swedish Ethical Review Board and from the Steering Committee of the Swedish Twin Registry. Researchers using the data are required to follow the terms of an agreement containing a number of clauses designed to ensure protection of privacy and compliance with relevant laws. For further information, contact Patrik Magnusson.

## Declaration of interests

The authors state the following competing interests: MU is a cofounder of Atlas Antibodies AB, which sells HPA antibodies used in this study. BF, FE, LF, JMS acknowledge a relationship with Atlas Antibodies AB. The other authors declare that they have no competing interests.

## Acknowledgements

We like to thank all participants of the S3WP study and the staff nurses collecting the samples. We acknowledge The Swedish Twin Registry for access to samples and data. We also thank every current or former member of the Affinity Proteomics Division at SciLifeLab, especially Elin Birgersson, Philippa Pettingill, Sofia Bergström, and Cecilia Mattson for technical support. We thank Helian Vunk for targeted proteomics data and Valtteri Wirta’s team for their sequencing work. We also thank the entire staff of the Human Protein Atlas for their tremendous efforts.

## Author Contributions

TDC, RSH, AB, MD, FE, BF performed experiments. TDC, MGH, CET, FE and JMS performed data analysis. PKEM provided TwinGene samples and supervised related analyses. LF and ISK managed the S3WP project samples and data. JO provided clinical expertise. AG and GB coordinated the collection of S3WP samples and clinical data. GB and MU provided funding and supervised the S3WP study. TDC and JMS conceived the experiments and wrote the manuscript with input from all co-authors. All co-authors approved the final version of the manuscript.

